# Evolution of a fatty acyl-CoA elongase underlies desert adaptation in *Drosophila*

**DOI:** 10.1101/2023.02.04.527143

**Authors:** Zinan Wang, Jian Pu, Cole Richards, Elaina Giannetti, Haosu Cong, Zhenguo Lin, Henry Chung

## Abstract

To survive in extreme environments such as hot-arid deserts, desert-dwelling species have evolved physiological traits to withstand the high temperatures and low aridity beyond the physiologically tolerable ranges of most species. Such traits which include reducing water loss have independently evolved in multiple taxa. However, the genetic and evolutionary mechanisms underlying these traits have thus far not been elucidated. Here we show that *Drosophila mojavensis*, a fruitfly species endemic to the Sonoran and Mojave deserts, had evolved extremely high desiccation resistance, by producing very long chained methyl-branched cuticular hydrocarbons (mbCHCs) that contributes to a cuticular waterproofing lipid layer reducing water loss. We show that the ability to synthesize these longer mbCHCs is due to evolutionary changes in a fatty acyl-CoA elongase (*mElo*). CRISPR/Cas9 knockout of *mElo* in *D. mojavensis* led to loss of longer mbCHC production and significant reduction of desiccation resistance at high temperatures but did not affect mortality at high temperatures or desiccating conditions individually, indicating that this gene is crucial for desert adaptation. Phylogenetic analysis showed that *mElo* is a *Drosophila* specific gene with no clear ortholog outside Diptera. This suggests that while the physiological mechanisms underlying desert adaptation are general, the genetic mechanisms may be lineage-specific.

## INTRODUCTION

The divergence and evolution of adaptive traits allows organisms to survive and thrive in diverse and extreme environments (Bardgett and Van Der Putten, 2014; McDonnell and Hahs, 2015; Rahbek et al., 2019). A key feature of these extreme environments is having multiple abiotic factors of which levels are beyond the physiologically tolerable ranges of most species (Hofmann and Todgham, 2010). In many cases, the interaction between abiotic factors may exacerbate the stresses caused by these factors to organisms that live in the environments (Filazzola et al., 2021; Mittler, 2006; Zhang et al., 2022). For example, in hot-arid deserts, the increased organismal water loss due to high vapor pressure deficit caused by high levels of aridity is exacerbated by high temperatures leading to even more rapid water loss (Cloudsley-Thompson, 1975; Gibbs et al., 1998). Nevertheless, species that are able to survive in these environments had evolved traits that allow them to withstand these stresses.

To survive rapid water loss in deserts, species from different taxa evolved high levels of desiccation resistance *via* similar physiological changes such as reducing water evaporation from the body, lowering metabolism, and minimizing water excretion (Gibbs, 2002; Merkt and Taylor, 1994; Williams and Tieleman, 2005). While there are some research on these independently evolved physiological traits (Gibbs and Matzkin, 2001; Rocha et al., 2021a; Xu et al., 2020), the underlying molecular and evolutionary mechanisms remain largely unknown. Recent association studies using genomic and transcriptomic studies have identified some candidate genes that may contribute to physiological adaptation in desert organisms (Gonzalez-Tokman et al., 2020; Rocha et al., 2021b; Wang et al., 2021), but the function of these genes are not characterized. In addition, it is not clear whether these adaptive traits in diverse desert species share the underlying same genetic mechanisms or are specific to different lineages of species. Determining the genetic basis underlying the evolution of desert adaptative traits may allow the prediction of whether and how contemporary species will evolve and adapt to future environmental changes such as global desertification (Huang et al., 2016; Shi et al., 2021).

In this study, we investigated the genetic basis underlying desert adaptation in a widely studied desert species, *D. mojavensis* (Gibbs, 2002; Matzkin and Markow, 2009). This species has adapted to several non-habitable deserts in southern California and Mexico (Reed et al., 2007), such as the Sonoran Desert where the relative humidity in the summer can be lower than 10% and the air temperature routinely exceeds 40°C (Gibbs et al., 2003). *D. mojavensis* has one of the highest levels of desiccation resistance (Kellermann et al., 2012) and the lowest rate of water loss in desiccating environments among *Drosophila* species (Gibbs and Matzkin, 2001). We showed that the high desiccation resistance of *D. mojavensis* at these desert conditions is due to its ability to synthesize very long chained methyl-branched cuticular hydrocarbons (mbCHCs), which contributes to a waterproofing lipid layer on its cuticle, reducing water loss. The synthesis of these very long chained mbCHCs is due to coding differences in a fatty acyl-CoA elongase (*mElo*) that allows *D. mojavensis* to synthesize longer mbCHCs compared to *D. melanogaster*, a well-studied cosmopolitan species. Phylogenetic analyses showed that *mElo* is a lineage-specific gene in *Drosophila* and two sibling genera in the same subfamily, suggesting that the evolution of *mElo* may contribute to the adaptation to future warmer and drier environments in species of these genera.

## RESULTS

### The fatty acyl-CoA elongase *mElo* (*CG18609*) elongates methyl-branched CHCs (mbCHCs) in *D. melanogaster*

We had previously shown that the length of mbCHCs can largely explain desiccation resistance across *Drosophila* species (Wang et al., 2022). *Drosophila* species produces combinations of mbCHCs of different carbon backbone lengths ranging from 24 carbons (2MeC24) to 32 carbons (2MeC32) (Jallon and David, 1987; Khallaf et al., 2021; Wang *et al*., 2022). *D. melanogaster* mainly produces 2MeC24, 2MeC26, and 2MeC28, while *D. mojavensis* produces longer mbCHCs, 2MeC28, 2MeC30, and 2MeC32 (**Figure 1A**). As CHCs are synthesized via the fatty acyl-CoA synthesis pathway, before decarbonylation to hydrocarbons in insect oenocytes, we hypothesized that an oenocyte specific fatty acyl-CoA elongase may underlie differences in the mbCHC chain lengths between these two species (Blomquist and Ginzel, 2021; Chung and Carroll, 2015; Holze et al., 2021). A previous genome wide association study in *D. melanogaster* showed that RNAi of a specific fatty acyl-CoA elongase, *CG18609*, reduces the production of mbCHCs (Dembeck et al., 2015). To confirm the role of *CG18609* in the elongation of mbCHCs, we used CRISPR/Cas9 to knock out this gene in *D. melanogaster*. While homozygous *CG18609* knockout strains are viable and fertile, levels of 2MeC26 and 2MeC28 were significantly reduced (**Figure 1B**). Oenocyte-specific GAL4/UAS expression of a *D. melanogaster CG18609* transgene in the homozygous knockout strain was able to restore production of 2MeC26 and 2MeC28 (**Figure 1B**). This suggests that *CG18609* is an elongase gene that is involved in the synthesis pathway of the fatty acyl-CoA precursors for 2MeC26 and 2MeC28 (**Figure 1C**). We named this gene *mElo* (***m****bCHC* ***Elo****ngase*).

**Figure 1.**
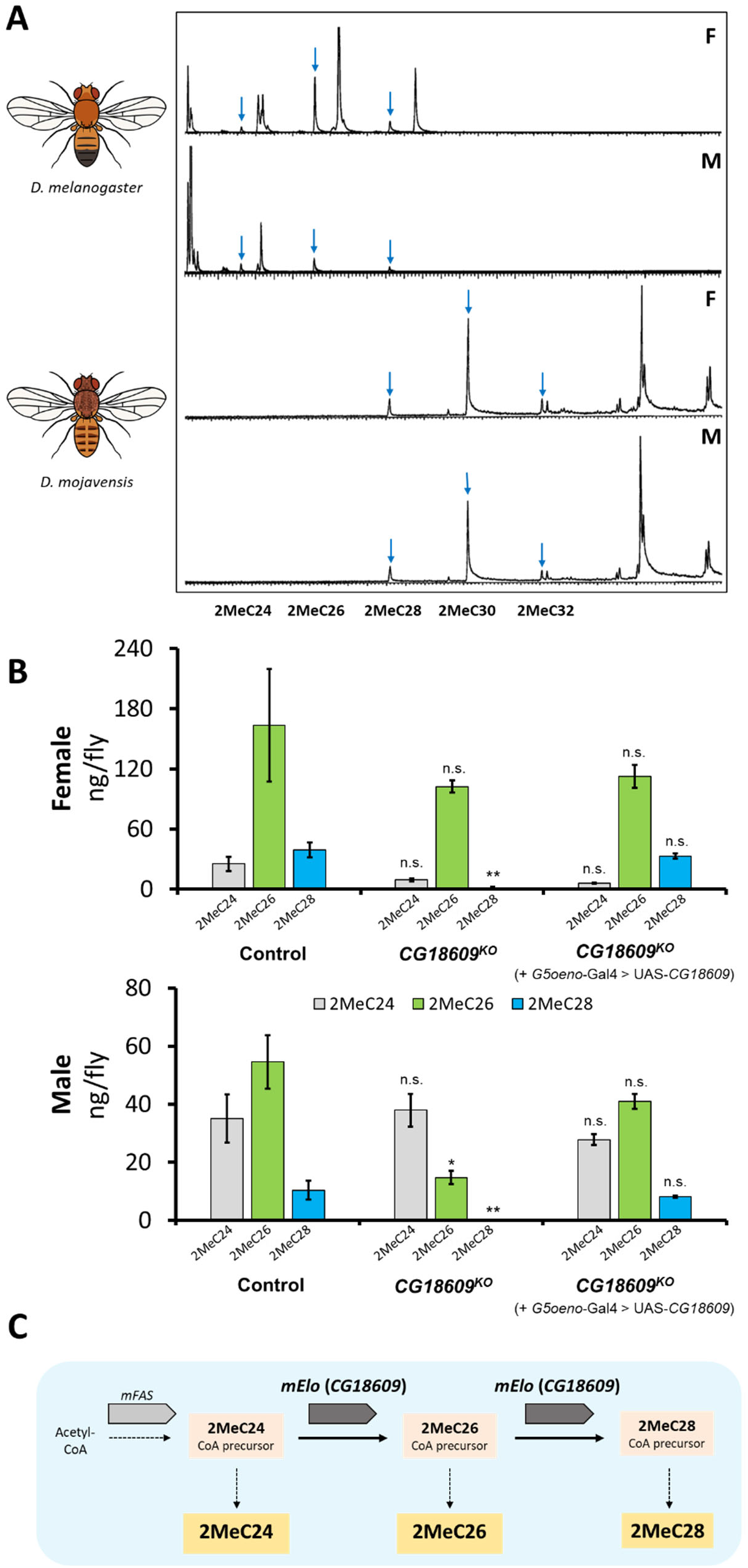
*mElo* (*CG18609*) is a methyl-branched CHCs (mbCHCs) *elongase* in *D. melanogaster*. **(A)** Gas chromatography mass spectrometry (GC-MS) chromatograms showing male and female CHCs of *D. melanogaster* and *D. mojavensis*. The desert *Drosophila* species *D. mojavensis* produces longer mbCHCs than the cosmopolitan *D. melanogaster*. Blue arrows indicate mbCHCs in the chromatogram. **(B)** Levels of mbCHCs in *D. melanogaster CG18609* homozygous knockout and rescue strains with oenocyte specific expression of *CG18609* (*G5Oeno*-Gal4>UAS-*CG18609*) compared to the control strain *Cas9onIII*, which the knockout was derived from. In both sexes, the levels of 2MeC28 were significantly reduced (Welch t-test. Female: *t*_(4.1)_ = 4.9, *P* = 0.007; Male: *t*_(4)_ = 3.2, *P* = 0.03), while the level of 2MeC26 was only significantly reduced in males (*t*_(4.5)_ = 4.2, *P* = 0.01). No significant differences were observed in 2MeC24 in both sexes (Female: *P* = 0.09; Male: *P* = 0.8). The rescue strains were able to restore the production of mbCHCs in both sexes, leading to the same mbCHC profiles as shown in the control strain (2MeC24, Female: *P* = 0.06; Male: *P* = 0.4; 2MeC26, Female: *P* = 0.4; Male: *P* = 0.2; 2MeC28, Female: *P* = 0.5; Male: *P* = 0.5). **(C)** The role of *CG18609* (*mElo*) in the elongation of 2MeC24 to 2MeC26 and 2MeC28 in *D. melanogaster*, based on knockout and rescue data.

### Transgenic overexpression of the *D. mojavensis mElo* (*Dmoj/mElo*) gene *in D. melanogaster* leads to longer mbCHC production and higher desiccation resistance

To investigate the molecular mechanisms underlying longer mbCHCs in *D. mojavensis*, we focused on the *mElo* gene of this species. At the *D. melanogaster mElo* locus, there are two elongase genes, *mElo* and another elongase gene, *CG17821*, while the *mElo* locus in *D. mojavensis* contains four predicted elongase genes, *GI20343, GI20344, GI20345*, and *GI20347* (**Figure 2A**). Phylogenetic analyses suggest that *GI20347* is the ortholog of *mElo*, while *CG17821* is likely to be orthologous with *GI20343, GI20344* and *GI20345* (**Figure S1**). We named *GI20347* as *Dmoj/mElo. In situ* hybridization with antisense probes of these genes showed that *mElo* is expressed in adult *D. melanogaster* oenocytes, while *GI20343, GI20345*, and *GI20347* (*Dmoj/mElo*) are expressed in adult *D. mojavensis* oenocytes. *CG17821* and *GI20344* are not expressed in *D. melanogaster* and *D. mojavensis* oenocytes respectively (**Figure S2**).

**Figure 2.**
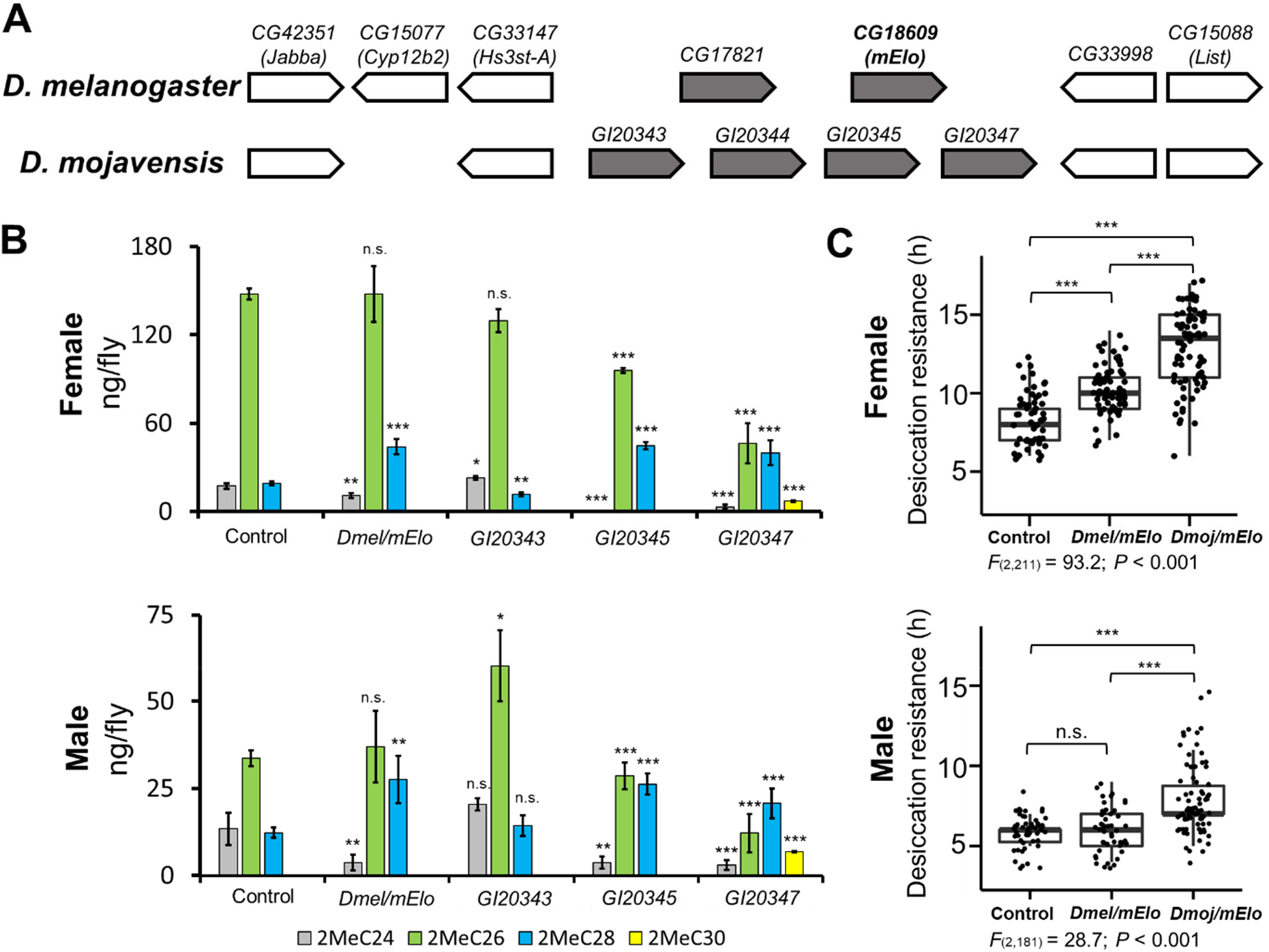
Oenocyte overexpression of *GI20347* in *D. melanogaster* leads to production of longer mbCHCs and confers higher desiccation resistance. **(A)** Microsynteny at the *mElo* locus is conserved between *D. melanogaster* and *D. mojavensis*. In *D. melanogaster*, two elongase genes, *CG17821* and *mElo* are present at this locus. In *D. mojavensis*, four elongase genes (*GI20343, GI20344, GI20345*, and *GI20347*) are present. **(B)** Quantities of mbCHCs (ng/fly) in *D. melanogaster* with each of the elongase genes (*mElo, GI20343, GI20345*, and *GI20347*) overexpressed in adult oenocytes using an oenocyte specific driver. The quantity of each mbCHC in the overexpression strains was compared with the control to determine any significant differences using the student’s *t*-test at *alpha*=0.05. **(C)** Desiccation resistance of *D. melanogaster* strains with *mElo* and *GI20347* (*Dmoj/mElo*) overexpressed in the oenocytes. Desiccation resistance is measured in hours (h) to mortality in a desiccating environment. Experiments were performed at 27°C for GAL4/UAS. Overexpression of *Dmoj/mElo* in *D. melanogaster* confers significantly higher desiccation resistance in both males and females compared to control strains or strains with overexpression of *Dmel/mElo*. One-way ANOVA was used to determine the differences in desiccation resistance between the strains of *D. melanogaster*, following with *post hoc* comparisons using Tukey’s method. *: *P* < 0.05; **: *P* < 0.01; ***: *P* < 0.001.

To determine the function of these elongase genes in mbCHC production, we overexpressed *mElo, GI20343, GI20345*, and *GI20347* individually in adult *D. melanogaster* oenocytes using the GAL4/UAS system at 27°C. Overexpression of *GI20343* did not change mbCHC production in males but led to slightly reduced 2MeC28 and increased 2MeC24 in females (**Figure 2B; Table S1**). The overexpression of *mElo* in *D. melanogaster* led to an increase in 2MeC28 production and a decrease in 2MeC24 production, which is similar to what we have shown in *mElo* homozygous knockout flies (**Figure 1B**). Overexpression of *GI20343* and *GI20345* individually in the oenocytes altered proportions of 2MeC24, 2MeC26, and 2MeC28 produced but did not result in the production of any longer mbCHCs. However, when we overexpressed *GI20347* (*Dmoj/mElo*), we observed a shift to the production of longer CHCs, including the increased production of a longer mbCHC, 2MeC30, which is usually absent or present in trace amounts in *D. melanogaster* (**Figure 2B**). As *GI20347* is the *D. mojavensis* ortholog of *D. melanogaster mElo*, we suggest that protein coding differences in this elongase gene may underlie the differences in mbCHC production between these two *Drosophila* species.

These overexpression strains allowed us to test our hypothesis that the production of longer mbCHCs may confer higher desiccation resistance in *Drosophila* species, allowing species to survive in desert conditions. To test this, we performed desiccation assays on the strains with *Dmel/mElo* and *Dmoj/mElo* overexpression. Our experiments showed that transgenic *D. melanogaster* flies with *Dmoj/mElo* overexpression were significantly more desiccation resistant (Mean ± SE, Females: 13.0 ± 0.3 h, Males: 7.8 ± 0.2 h) compared to control flies (Females: 8.4 ± 0.2 h, Males: 5.9 ± 0.1 h) and flies with *Dmel/mElo* overexpression (Females: 10.3 ± 0.2 h, Males: 6.0 ± 0.2 h) (**Figure 2C**). This result demonstrated that the production of longer mbCHCs can significantly increase desiccation resistance, consistent with our previous findings using synthetic mbCHCs (Wang *et al*., 2022).

### *D. mojavensis mElo (Dmoj/mElo)* confers high desiccation resistance at desert temperatures

While our experiments showed that transgenic overexpression of *Dmoj/mElo* in *D. melanogaster* produces longer mbCHCs such as 2MeC30 and confers significantly higher desiccation resistance, we did not recapitulate the production of 2MeC32 and the very high desiccation resistance in the desert dwelling *D. mojavensis* (Wang *et al*., 2022). To investigate the role of *Dmoj/mElo* in mbCHC synthesis and desiccation resistance in *D. mojavensis*, we used CRISPR/Cas9 to knockout *Dmoj/mElo* in *D. mojavensis*. Three independent *Dmoj/mElo* knockout strains, M3.5, M3.9, and M3.11, carrying a 5-bp insertion, 90-bp deletion, and 10-bp deletion in the exon 3 of *Dmoj/mElo*, respectively, were obtained (**Figure 3A, Figure S3**). All three mutant strains are homozygous viable. We also established three independent isofemale strains, ISO1, ISO2, and ISO3, from the parental population as controls.

**Figure 3.**
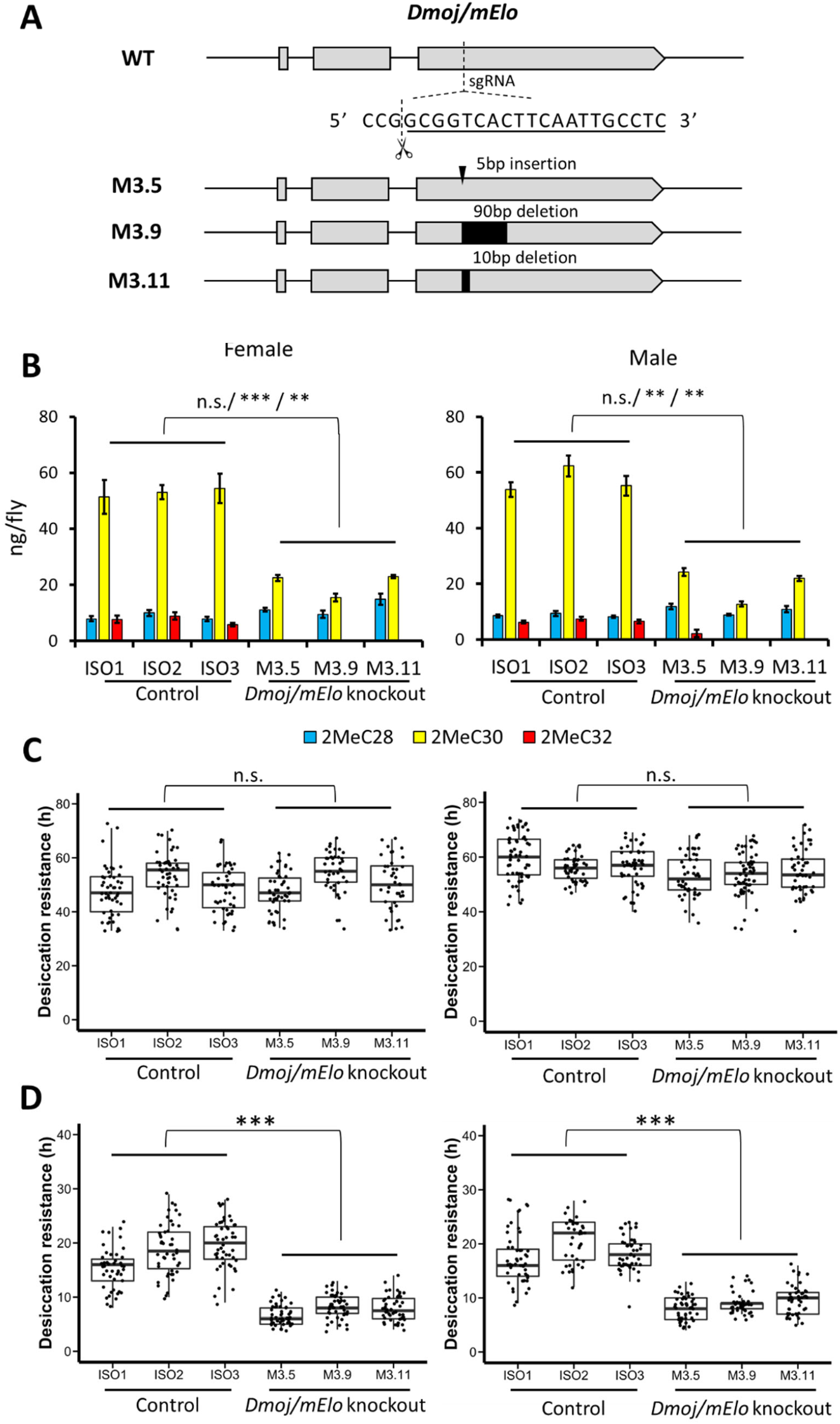
Knockout of the *mElo* ortholog *GI20347* (*Dmoj/mElo*) in *D. mojavensis* leads to a significant decrease in desiccation resistance at an ecologically relevant high temperature. **(A)** A CRISPR/Cas9 non-homologous end-joining strategy resulted in three homozygous viable strains with *Dmoj/mElo* knockout, M3.5, M3.9 and M3.11 in *D. mojavensis*, which have a 5 bp insertion, a 90 bp deletion, and a 10 bp deletion, respectively. **(B)** In the three *Dmoj/mElo* (*GI20347*) knockout strains, 2MeC30 was significantly reduced (∼50% of wild type levels) and 2MeC32 is reduced to trace amounts (Female: 2MeC30: *t*_(4)_ = -11.6, *P* < 0.001, 2MeC32: *t*_(4)_ = -8.5, *P* = 0.001; Male: 2MeC30: *t*_(4)_ = -8.5, *P* = 0.001, 2MeC32: *t*_(4)_ = -7.3, *P* = 0.002). **(C)** There are no significant differences in desiccation resistance between the three *Dmoj/mElo* knockout strains and the three isofemale control strains at 27°C (Female: *P* = 0.7; Male: *P* = 0.06). **(D)** At 37°C, the three *Dmoj/mElo* knockout strains have a significant reduction in desiccation resistance compared to the three isofemale control strains (Female: *t*_(4)_ = 7.4, *P =* 0.002; Male: *t*_(4)_ = 9.5, *P* < 0.001). For both CHC quantities and desiccation resistance, linear mixed effects models were applied to compare the two groups of flies using ‘*lmer*’ function in R (ver 4.1). The three isofemale wild type and independent knockout strains were included as random effects. The difference between the wildtype and knockout flies was determined by paired contrast at *alpha* = 0.05.

In all three *Dmoj/mElo* knockout strains, 2MeC32, the longest mbCHC in *D. mojavensis*, was reduced to trace amounts and 2MeC30 was significantly reduced compared to the control strains (**Figure 3B, Figure S4, Table S2**), suggesting that *Dmoj/mElo* is responsible for elongating 2MeC28 to 2MeC30 and 2MeC32 in *D. mojavensis*. We further examined how these changes in mbCHC lengths could affect desiccation resistance of *D. mojavensis* by subjecting all knockout and control strains to the desiccation assay at 27°C. However, we did not observe any significant difference in desiccation resistance between the knockout strains and the controls at this temperature (**Figure 3C**). As the capability of CHCs in preventing water loss is associated with their melting temperatures (Gibbs, 2007; Wigglesworth, 1945), and the air temperature of the microhabitat of *D. mojavensis* (e.g., outside cactus necrosis in the Sonoran Desert) is higher than 35 °C (Gibbs *et al*., 2003), we considered the hypothesis that at these higher temperatures, longer mbCHCs such as 2MeC32 may make a difference in desiccation resistance. Therefore, we tested whether the reduced 2MeC30 and 2MeC32 in *Dmoj/mElo*-knockout *D. mojavensis* could affect its desiccation resistance at 37°C, a temperature that is ecologically relevant to *D. mojavensis*.

Desiccation experiments at 37°C showed that across the board, time to mortality is faster than experiments performed at 27°C. At this temperature, the three *Dmoj/mElo* knockout strains are significantly less desiccation resistant (Females: 7.7 ± 0.2 h, Males: 8.9 ± 0.2 h) compared to the control strains (Females: 18.0 ± 0.4 h, Males: 18.6 ± 0.4 h) (**Figure 3D**), suggesting that the production of longer mbCHCs such as 2MeC30 and 2MeC32 is crucial in desiccation resistance in hot and dry conditions. To exclude the possibility that the higher mortality of the *Dmoj/mElo* knockout flies compared to the control flies was due to heat stress rather than increased water loss at 37°C, we tested the survival of adults of both knockout and control strains at 37°C under a non-desiccating experimental environment (flies are given fresh food every 2-3 days). Survival at 37°C between the control strains and the *Dmoj/mElo* knockout strains were not significantly different under these conditions (**Figure S5**), suggesting that the increased mortality observed during the desiccation experiment at 37°C was due to water loss rather than the higher temperature. Taken together, our results demonstrated that in *D. mojavensis, Dmoj/mElo* underlies the production of long mbCHCs such as 2MeC30 and 2MeC32 and contributes to the high desiccation resistance of this species in its hot and arid desert environment.

### The *mElo* gene is a *Drosophila* specific mbCHC elongase

As mbCHCs are almost ubiquitous in most insect species, we sought to investigate if the role of *Dmoj/mElo* in determining mbCHC length and desiccation resistance is conserved across Insecta. Using the conserved microsynteny (*Jabba, Cyp12b2, Hs3st-A, CG33998*, and *List*) around *CG17821* and *mElo* between *D. melanogaster* and *D. mojavensis*, we investigated this locus 16 *Drosophila* species and five species from closely related genera (*Scaptodrosophila, Chymomyza, Leucophenga, Phortica*, and *Ephydra*) (Kim et al., 2021; Scott et al., 2014; Vicoso and Bachtrog, 2015) (**Figure 4A**). We found that in the *Drosophila* species examined, this microsynteny is conserved and the elongase gene copy number ranges from two to four across these *Drosophila* species. This microsynteny is also conserved in *Scaptodrosophila lebanonensis, Chymomyza costata*, and *Leucophenga varia*, and partially conserved in *Phortica variegata, Ephydra gracilis*, and *Musca domestica*. There are two elongase genes at this locus in *S. lebanonensis* and *C. costata*, but none in *L. varia, P. variegata*, and *E. gracilis*. This suggests that elongase genes at this locus first originate in the common ancestor of the *Drosophila, Scaptodrosophila*, and *Chymomyza* genus (i.e., the *Drosophilinae* subfamily) (**Figure 4A**).

**Figure 4.**
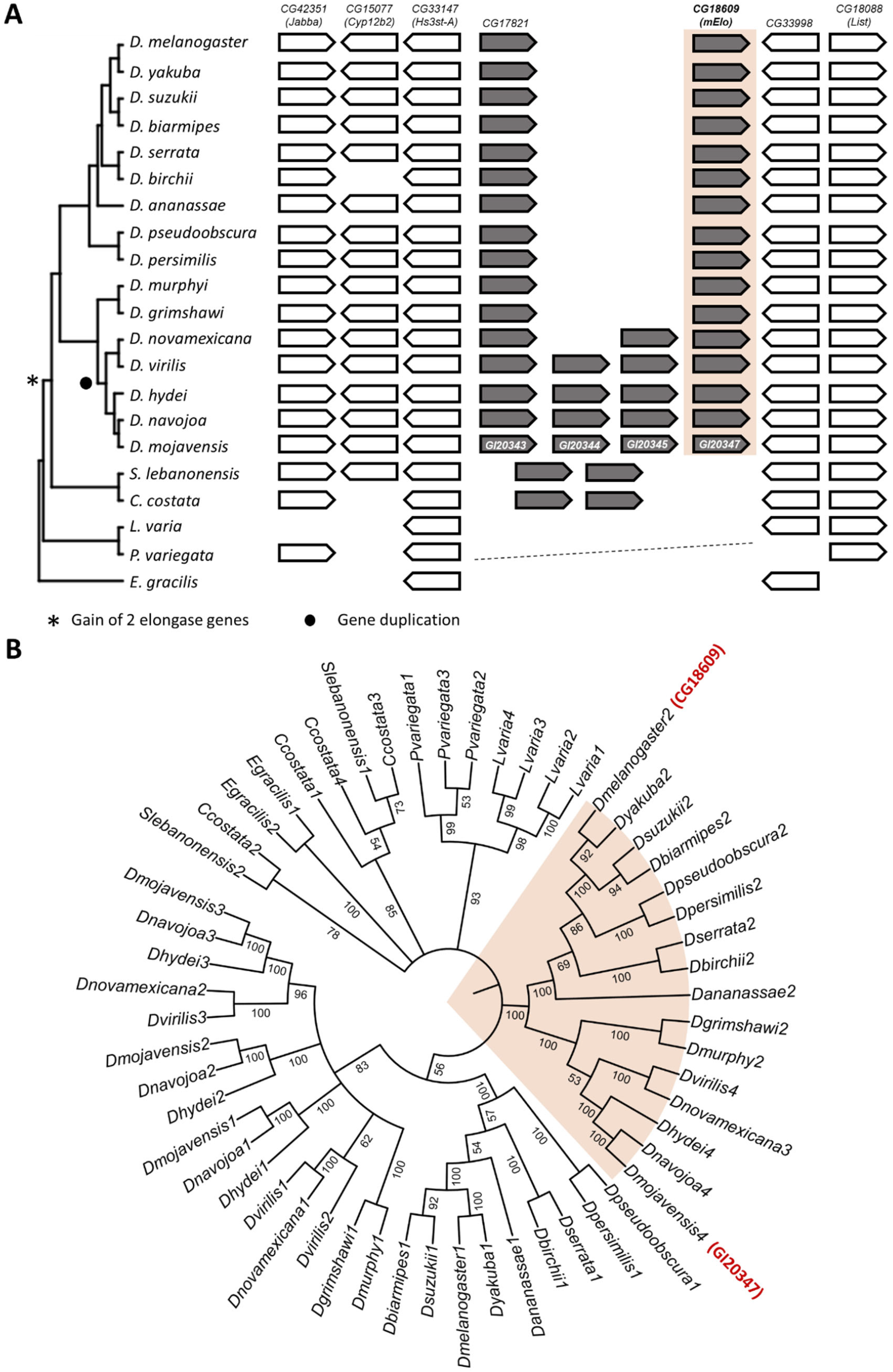
The origins of *mElo* in *Drosophila*. **(A)** The *mElo* loci in 16 *Drosophila* species and species from six closely related genera. The *mElo* loci were identified based on the conserved genes in the *D. melanogaster mElo* locus (*Jabba, Cyp12b2, Hs3st-A, CG33998*, and *List*) which are used as anchor genes in our analysis. All *Drosophila* species contains at least two elongase genes at this locus. There was an expansion of elongase genes in the virilis and repleta clades where species have 3-4 elongase genes at this locus (denoted by a *). Two elongase genes are present in *S. lebanonensis* and *C. costata*, but none in *L. varia, P. variegata*, and *E. gracilis*. This suggests that elongase genes at this locus first originated in the common ancestor of the *Drosophila, Scaptodrosophila*, and *Chymomyza* genus (denoted by a *). The dashed line indicates that anchor genes can be located in the genome but at different location. **(B)**. Phylogenetic relationship of elongase genes in *mElo* loci of Drosophilinae species as well as the elongases from *S. lebanonensis, C. costata, L. varia, P. variegata*, and *E. gracilis* that share the highest similarities to *mElo*. The phylogenetic tree was inferred by the ML method using amino acid sequences with 1000 bootstrap tests. The numbers next to nodes represent bootstrap values.

To determine the relationship of these elongase genes, we performed a phylogenetic analysis of all these elongase genes at this locus from 16 *Drosophila* species, *S. lebanonensis*, and *C. costata*. We also included elongase genes in *L. varia, P. variegata*, and *E. gracilis* which has the highest sequence homology to the elongases at the *Drosophila mElo* locus. Phylogenetic analysis of all elongase genes in *mElo* loci showed that each *Drosophila* species only has a single *mElo* ortholog (**Figure 4B**). In addition, the elongase genes in *Scaptodrosophila* and *Chymomyza* did not cluster with those in *Drosophila* (**Figure 4B**), suggesting that the presence of multiple elongase genes in the two lineages is likely due to lineage-specific gene duplication events. This result suggests that the *mElo* gene at the *Drosophila mElo* locus originated in the genus *Drosophila*. However, this does not exclude the possibility that the *mElo* gene is present in other insect species, but located in another genomic location, as mbCHCs are prevalent across insect species. To determine if any *mElo* ortholog is present in other insect species, we compared elongase genes *Aedes aegypti*, a Dipteran mosquito species and several non-Diptera species, *Apis mellifera, Bombyx mori*, and *Tribolium castaneum*. From our phylogenetic analysis, we observed that while other elongase genes such as *sit* and *CG31523* have 1:1 ortholog in all insect species, there is no clear *mElo* orthologous gene identified (**Figure S6**). This suggests that *mElo* gene is a *Drosophila* specific mbCHC elongase and other elongase genes may elongate mbCHCs in other insect species.

## DISCUSSION

A few of the many species on Earth have evolved adaptive traits to live in extreme environments with harsh abiotic conditions. However, few studies have determined the underlying genetic basis for such traits. In this study, we show that the desert *Drosophila* species, *D. mojavensis*, has evolved coding changes in a fatty acyl-CoA elongase gene, *mElo*, which led to the production of very long mbCHCs and high desiccation resistance in this species. While the knockout of this gene in *D. mojavensis* has no significant effects on desiccation resistance at a lower temperature (27°C), it significantly reduces desiccation resistance at a higher temperature (37°C), which is within the average high temperature range in the Sonoran Desert during summer (**Figure S7**). This suggests that these very long mbCHCs are able to reduce water loss at hot-arid desert conditions i.e., high temperature and low humidity, and is crucial for survival in this habitat. The transgenic overexpression of the *D. mojavensis mElo* gene in the cosmopolitan *D. melanogaster* led to the production of longer mbCHCs and higher desiccation compared to the transgenic overexpression of the *D. melanogaster mElo* gene, suggesting evolved coding differences in this gene between the two species. Finally, phylogenetic analyses of this locus suggest that the *mElo* gene evolved recently and an orthologous copy of this gene is not found outside Diptera.

### The very long mbCHCs in *D. mojavensis* are critical for survival in hot and arid deserts

Why are there significant differences in desiccation resistance at 37°C but not at 27°C between *Dmoj/mElo* knockout strains and the control strains? The ability of the CHC layer to prevent water loss depends on the physical state of this solid-liquid mixture layer and this affects its ability to prevent water molecules from diffusing through (Menzel et al., 2019). At a specific “phase transition” temperature, water loss through the cuticle increases rapidly (Gibbs and Pomonis, 1995; Wigglesworth, 1945). This transition temperature is determined by the CHC composition of each species (Gibbs and Pomonis, 1995; Menzel *et al*., 2019).

We suggest that the loss of 2MeC32 and the significant decrease of 2MeC30 in the *Dmoj/mElo* knockout strains altered the transition temperature of the CHC layer on *D. mojavensis*. At 27°C, this does not affect the *Dmoj/mElo* knockout strains, therefore they do not differ significantly in desiccation resistance from the control strains. However, at 37°C, *Dmoj/mElo* knockout strains begin to lose water more rapidly than the control strains, resulting in a significant decrease in desiccation resistance compared to the control strains (**Figure 5B**). As hot and arid deserts have long days of high temperatures during the summer, we suggest that the very long mbCHCs in *D. mojavensis* are crucial for survival as they allow *D. mojavensis* to survive the hot and dehydrating conditions during the long day before the dip in temperatures during the night.

**Figure 5.**
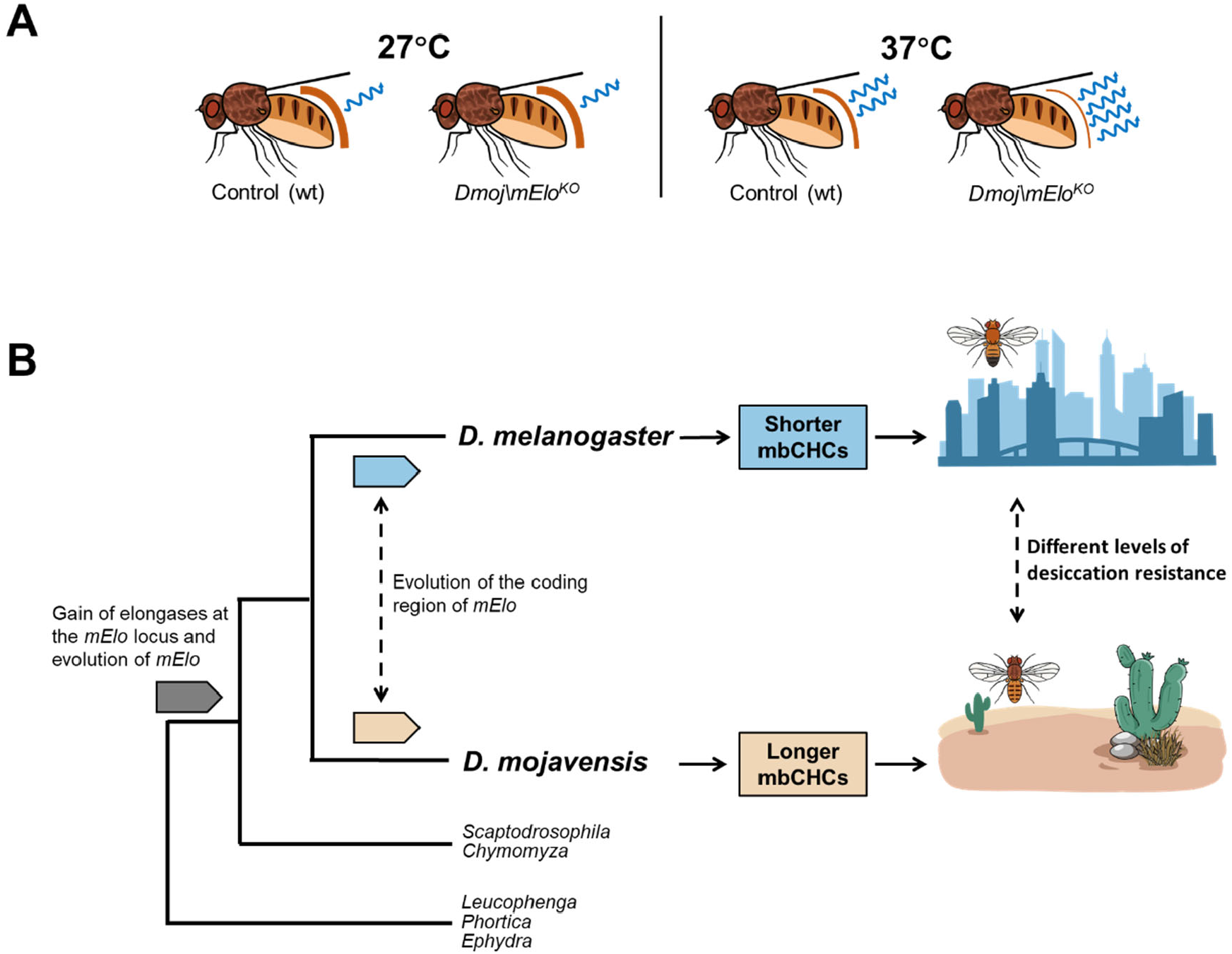
Evolution a fatty acyl-CoA elongase gene, *mElo*, underlie higher desiccation resistance and desert adaptation in *D. mojavensis*. **(A)** A schematic showing that the loss of 2MeC32 and a significant amount of 2MeC30 at 27°C does not affect the *Dmoj/mElo* strain of *D. mojavensis* compared to the control strain as water loss is similar between these strains. However, at the higher temperature of 37°C, we hypothesized the *Dmoj/mElo* strain loses water more rapidly compared to the control strain due to the melting temperatures of the CHC layer being altered by the loss of these longer mbCHCs. **(B)** A model showing how coding changes in *mElo* led to shorter mbCHCs in the cosmopolitan species *D. melanogaster* and longer mbCHCs in the desert species, *D. mojavensis*, allowing it to survive in the hot and dry desert.

### Evolution at the *mElo* locus in *Drosophila*

The oenocyte driven overexpression of *Dmel/mElo* and *Dmoj/mElo* in the *D. melanogaster* produces mbCHCs of different chain lengths (**Figure 2**), suggesting that there are differences in protein coding sequences of this gene between the two *Drosophila* species, and that these differences contribute to the different mbCHCs produced by these two species. Our previous study using ancestral trait reconstruction showed that the last common ancestor of *D. melanogaster* and *D. mojavensis* has an mbCHC phenotype that is intermediate between the mbCHCs phenotypes of the two species (Wang *et al*., 2022). As *mElo* controls the length of the longest mbCHCs produced in each of these two species, this suggests that evolutionary changes in this gene may have occurred in both species from the common ancestor, i.e., evolution in this gene led to *D. melanogaster* to produce shorter mbCHCs and *D. mojavensis* to produce longer mbCHCs as both species adapt to their environments (**Figure 5B**).

The CRISPR/Cas9 knockout of *mElo* in both species also produced different mbCHC phenotypes. In *D. melanogaster*, knockout of *mElo* produced a mbCHC phenotype which is largely 2MeC24 in males and 2MeC26 in females with significant decreases in 2MeC28 in both sexes compared to wild-type flies. In *mElo* knockout *D. mojavensis*, while 2MeC30 and 2MeC32 are significantly reduced with the latter reduced to trace amounts compared to wild-type flies, the major CHC in these *mElo* knockout *D. mojavensis* is still 2MeC30. This suggests that there are other elongase genes contributing to the mbCHC phenotype in *D. mojavensis*. A candidate gene for this would be *GI20345*, another elongase gene in the *mElo* locus in *D. mojavensis* that is expressed in *D. mojavensis* oenocytes and is able to elongate mbCHCs in *Dmel/mElo* knockout *D. melanogaster* (**Figure 2**). This may suggest a complicated evolutionary scenario in the evolution of mbCHC biosynthesis controlled by the *mElo* locus in *Drosophila* (**Figure S8**).

### Lineage specific genetic basis for the evolution of desiccation resistance

Variations in CHC composition contribute to desiccation resistance differences across many insect species (Buellesbach et al., 2018; Leeson et al., 2020; Rouault et al., 2004). While mbCHCs are found in almost all insect species, our phylogenetic analyses showed that the *mElo* gene is a *Drosophila* specific mbCHC elongase, indicating that the control of mbCHCs chain length in other species is likely to be a different elongase gene. This suggests that contribution to the evolution of desiccation resistance and desert adaptation by the *mElo* locus is likely to be lineage specific. If changes in CHC composition can contribute to desiccation resistance in insects, what are the likely genetic mechanisms that underlie the evolution of desiccation resistance beyond *Drosophila*? The CHC biosynthesis pathway is largely conserved in insects and is made up of several fatty acyl-CoA synthesis gene families such as fatty acyl-CoA synthetases, desaturases, fatty acyl-CoA reductases (FARs), and elongases (Blomquist and Ginzel, 2021). These gene families evolved rapidly and contribute to the diversification of CHCs across insects (Finck et al., 2016; Finet et al., 2019; Helmkampf et al., 2015; Tupec et al., 2019). Gains and losses of these genes as well as changes in their oenocyte expression are likely to contribute to CHC changes and the evolution of desiccation resistance in different insect species. The rapid “birth-and-death” of these genes also suggests that many of the genetic mechanisms leading to CHC changes and the evolution of desiccation resistance across different insect species are likely to be lineage specific.

## Conclusions

In summary, we showed that evolutionary change in a fatty acyl-CoA elongase contributes to the adaptation of *D. mojavensis* to the hot and arid Sonoran Desert by reducing water loss at a high temperature. While the general mechanisms of CHC composition modification leading to reduction of water loss at high temperatures in insects adapting to hotter and drier conditions are likely to be conserved, the specific genetic mechanism is not. This may have implications in the prediction of species changes as climate change continues to occur in the near future.

## ACKNOWLEGDEMENTS

We thank Ye Ma, Mei Luo, and Taylor Hori for technical assistance, and Yuzhang Shan for assistance with figure visualization. We acknowledge the Bloomington Drosophila Stock Center and National Drosophila Species Stock Center for fly stocks. This work is supported by a National Science Foundation grant (2054773) to H. Chung.

## AUTHOR CONTRIBUTIONS

Z.W. and H. Chung designed research; Z.W., J.P., H. Cong, E.G., C.R., Z.L., and H. Chung performed research; Z.W., Z.L., and H. Chung analyzed data; and Z.W. and H. Chung wrote the paper with input from other authors.

## MATERIALS AND METHODS

### *Drosophila* strains

The *y w; attP40* strain was used for *in situ* hybridization and transgenesis in *D. melanogaster*. The *D. mojavensis wrigleyi* strain (15081-1352.29) used was obtained from the National Drosophila Species Stock Center (NDSSC). The *oeno*GAL4 strain (*PromE(800) line 2M*) was a gift from Joel Levine (Billeter et al., 2009). The balancer strain *w*^*1118*^; *CyO/Sco; MKRS/TM6B, Tb*^*1*^ (#3703) and *y*^*1*^ *w*^*\**^ *P{y*^*+t7*.*7*^*=nos-phiC31\int*.*NLS}X; CyO/Sco* (#34770) were obtained from the Bloomington *Drosophila* Stock Center. All flies were maintained at room temperature on standard *Drosophila* food (Bloomington formulation, Genesee Scientific). *D. melanogaster* GAL4/UAS experiments were performed at 27^º^C.

### *In situ* hybridization & Imaging

*In situ* hybridization was performed on oenocytes of five-day-old adults using methods as described previously (Chung et al., 2007; Pu et al., 2021). Primers that were used for synthesizing probes were listed in **Table S4**. All *in situ* hybridization images were captured using the Nikon SMZ18 dissecting stereo microscope system.

### Generation of *mElo* knockout by CRISPR/Cas9 genome engineering in *D. melanogaster*

CRISPR/Cas9-mediated homology-directed repair (HDR) was used to generate a knockout of *Dmel/mElo*. The program, flyCRISPR Optimal Target Finder, was used to identify optimal CRISPR target sites (Gratz et al., 2014). Target-specific sequences for *Dmel/mElo* were synthesized as oligonucleotides, phosphorylated, annealed and ligated into the *BbsI* sires of *pU6-BbsI-chiRNA* (Addgene plasmid #45946) (Gratz et al., 2013) (5’: Dmel/mElo-gRNA1-BbsI-F and Dmel/mElo-gRNA1-BbsI-R, 3’: Dmel/mElo-gRNA2-BbsI-F and Dmel/mElo-gRNA2-BbsI-R). To construct the replacement donor, approximately 1kb homology arms flanking the cut sites were amplified by PCR using primers Dmel/mElo-RightHomo-AscI-F and Dmel/mElo-RightHomo-XhoI-R for the 5’ homology arm and primers Dmel/mElO-LeftHomo-EcoRI-F and Dmel/mElO-LeftHomo-NotI-R for the 3’ homology. The replacement donors were cloned sequentially into the corresponding cut sites of the dsDNA donor vector *pHD-DsRed-attP* (Addgene plasmid #51019). The primers used for generating gRNA and replacement donor constructs are listed in **Table S4**. The two gRNA constructs and the replacement donor construct were co-injected into the w^1118^; PBac{y^+mDint2^ GFP^E.3xP3^=vas-Cas9}VK00027 strain (denoted as *Cas9onIII* strain; BDSC #51324), which carries a *vasa-Cas9* transgene on Chromosome 3. The dsRed fluorescence in the eyes was used to screen positive progeny, which were then crossed to *w*^*1118*^ to remove the *vasa-Cas9* transgene before being back-crossed for five generations and then made homozygous using the double balancer strain *w*^*1118*^; *CyO/Sco; MKRS/TM6B, Tb*^*1*^. The replacement of *Dmel/mElo* with *attP*/*dsRED* by HDR was confirmed by PCR using the primers DmelCG18609-EcoRI-F and DmelCG18609-XbaI-R and the presence of dsRed (**Figure S9A**). The resulting strain is designated as *w*^*1118*^; *CG18609*^*KO-dsRED-attP*^ (*mEloKO*). A transgene carrying a *PhiC31* integrase driven by a *nanos* enhancer was integrated into this strain by crossing it to *y*^*1*^,*w*,P{y*^*+t7*.*7*^*=nos-phiC31\int*.*NLS}X; Sco/CyO* (**Figure S9B**). The resulting strain is *w*^*1118*^, *P{y*^*+t7*.*7*^*=nos-phiC31\int*.*NLS}X; CG18609*^*KO-dsRED-attP*^ and named as the *mEloKOint* strain.

### Generation of plasmid constructs

Primers used for generating all constructs are listed in **Table S4**. UAS overexpression constructs were cloned in *PhiC-31* site-specific transformation vector, *pWalium10-MOE* (Ni et al., 2009). The genomic DNA of *Dmel/CG17821, Dmel/CG18609* (*Dmel/mElo*), *Dmoj/GI20343, Dmoj/GI20344, Dmoj/GI20345*, and *Dmoj/GI20347* (*Dmoj/mElo*) were amplified by PCR from genomic DNA of corresponding species and then cloned into *pWalium10-MOE* vector using the *NdeI, EcoRI*, or *XbaI* sites. The *G5-GAL4* construct by cloning the 5’ regulatory region of *Dmoj/GI20345* into the GFP reporter vector *pS3aG* via the *AscI* and *SbfI* sites (Williams et al., 2008). The GFP sequence was then cut out from this construct using *SpeI* and *SbfI* and replaced with a GAL4 sequence *pBPGUw* (Addgene plasmid #17575) vector using *SpeI* and *SbfI*.

### Drosophila transgenesis and overexpression experiments

Transgenesis in *D. melanogaster* (*y w; attP40* and *mEloKOint* strains) was performed using the *PhiC31* integrase system following standard *Drosophila* transgenesis protocols. To generate overexpress UAS strains, the overexpression constructs of elongase genes on *pWalium10-MOE* were individually injected into the *y w; attP40* strain. The *G5-*GAL4 construct and the overexpression construct of *Dmel/mElo* on *pWalium10-MOE* were individually injected into the *mEloKOint* strain for the rescue of *mElo* expression in *mElo* knockout *D. melanogaster*. All overexpression experiments were performed at 27°C by reciprocally crossing *oeno*GAL4 strain (*oeno*GAL4 or *G5-*GAL4) and the corresponding UAS overexpression strain.

### Generation of *mElo* knockout by CRISPR/Cas9 genome engineering in *D. mojavensis*

To generate *Dmoj/mElo* mutant alleles in *D. mojavensis*, we used a non-homologous end joining mediated strategy by injecting the mixture of Cas9 protein (#CP01; PNA Bio) and sgRNAs into the embryos of this species. Following the protocol in Khallaf et al. (2020), we co-injected two sgRNAs targeting Dmoj/white (Dmoj/white_sgRNAa and Dmoj/white_sgRNAb). *Dmoj/mElo* specific sgRNAs (Dmoj/mElo-sgRNAa and Dmoj/mElo-sgRNAb) were designed using the online tool CRISPR Design (Gratz *et al*., 2013) and two sgRNAs were selected. All sgRNAs were generated following the protocol in Kistler et al. (2015), with *in vitro* transcription using T7 Megascript Kit (Ambion) and purification using a MegaClear Kit (Ambion). Primers used for the synthesis of all sgRNAs were listed in **Table S4**. The final injection mixture is composed of 300 ng/μL Cas9 protein and four sgRNAs, each 75 ng/μL. To screen for the offspring of *D. mojavensis* carrying *Dmoj/mElo* mutant alleles, we used the T7E1 assay (NEB #E3321) to determine potential mutations for every single fly following the protocol in (Zhu et al., 2019). To eliminate potential off-targets from the gene knockout, all strains carrying mutations in *Dmoj/mElo* were backcrossed with the parental *D. mojavensis* strain for at least five generations before being made homozygous.

### Cuticular hydrocarbon extraction and analyses

CHC extraction, GC/MS analysis, CHC identification, and quantification were performed as described previously (Wang *et al*., 2022). The GC thermal program was set as follows: start from 100 °C, 5 °C/min to 200 °C, and 3 °C/min to 325 °C. For each sex in each reciprocal cross, three extractions were conducted as replicates and the results were pooled for further statistical analyses, so six replicates were performed for each cross.

### Desiccation assay

Desiccation assays were performed as described previously (Wang *et al*., 2022). Silica gel (S7500-1KG) was ordered from Sigma-Aldrich. For each genotype, six replicates were conducted, each three from each reciprocal cross.

### Bioinformatics

The sequences of all elongase genes used in this study were retrieved from the NCBI (http://www.ncbi.nlm.nih.gov) database, VectorBase (Giraldo-Calderón et al., 2015), and SilkDB (Consortium, 2008) via TBLASTN using *CG17821* and *CG18609* as queries (Suppl. Fasta File). The DNA or amino acid sequences were aligned with MUSCLE and manually adjusted for the phylogenetic reconstruction using the maximum likelihood method in MEGA (Version 11) (Kumar et al., 2018). The GTR model with a Gamma distribution was applied to reconstruct phylogeny using CDS of elongase genes, while the LG substitution matrix and a Gamma distribution with invariant sites (G+I) was applied using their amino acid sequences. All phylogenetic reconstruction analyses used 1000 bootstrap replicates to test the reliability of inferred trees. The phylogenetic relationship of *Drosophila* and related species used in this study was adapted from (Finet et al., 2021; Pu *et al*., 2021).

## SUPPLEMENTARY FIGURES

**Figure S1.**
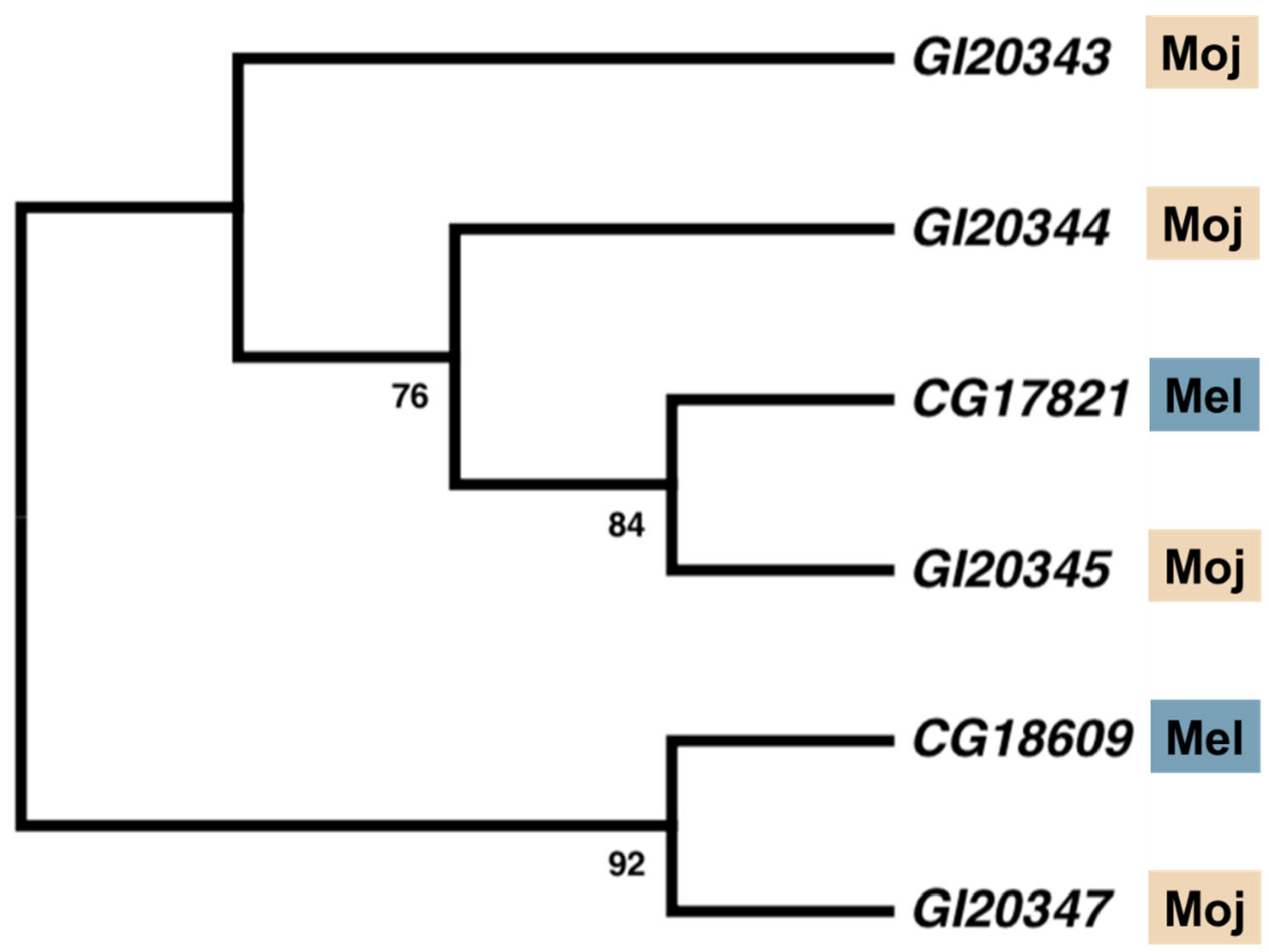
Phylogeny of elongases in the *mElo* loci of *D. melanogaster* and *D. mojavensis*. The coding sequences of these genes were used to generate the phylogeny using the Maximum Likelihood method with GTR model and 1000 bootstraps. The phylogenetic analysis showed that the *D. mojavensis* orthologue of *mElo* is *GI20347*.

**Figure S2.**
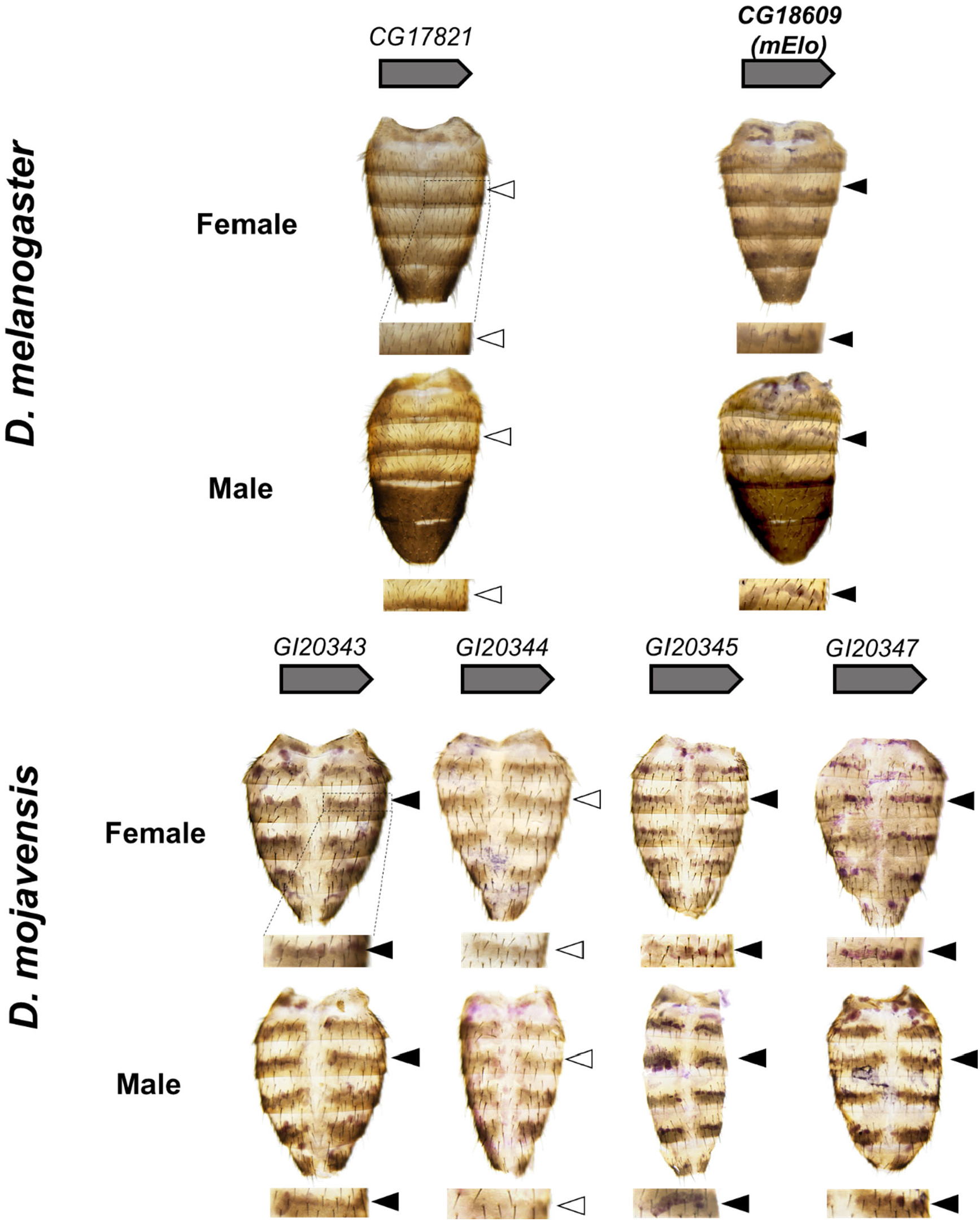
RNA *in situ* hybridization of fatty acyl-CoA elongase genes in the *D. melanogaster* and the *D. mojavensis mElo* loci on adults. In *D. melanogaster, CG18609* RNA transcript was detected in the adult oenocytes. In *D. mojavensis, GI20343, GI20345*, and *GI20347* RNA transcripts were detected in the adult oenocyte. The expression of all the four genes are sexually monomorphic. Arrowheads point to oenocytes. Filled arrowheads indicate visible expression detected and open arrowheads indicate no visible expression.

**Figure S3.**
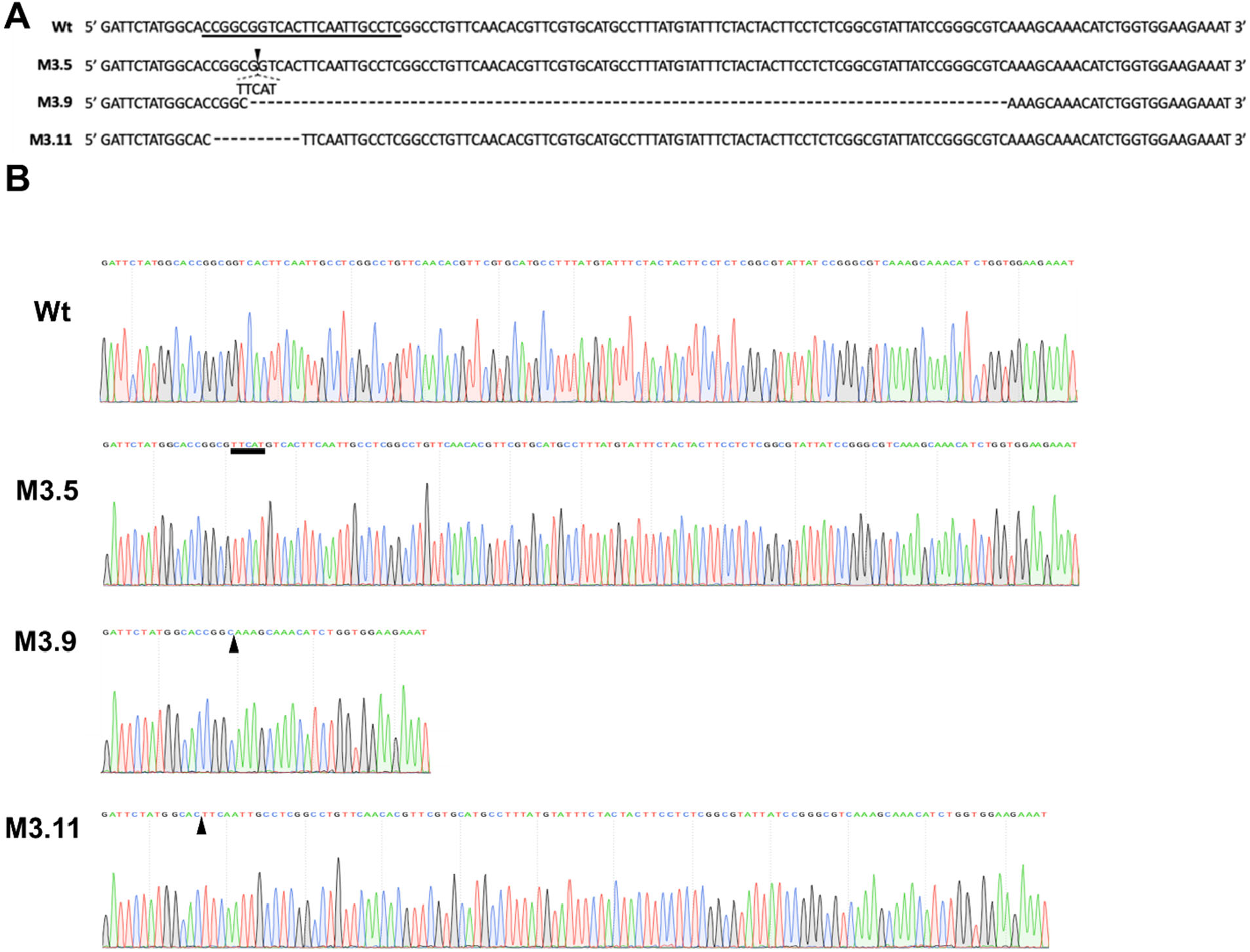
Edited sequences in M3.5, M3.9, and M3.11 strains. CRISPR/Cas9 and non-homologous end-joining was used to generate knockout strains of *GI20347*. Three independent knockout strains, namely M3.5, M3.9, and M3.11, were generated. They carry a 5-bp insertion, 90-bp deletion, and 10-bp deletion in the exon 3 of *Dmoj/mElo*, respectively.

**Figure S4.**
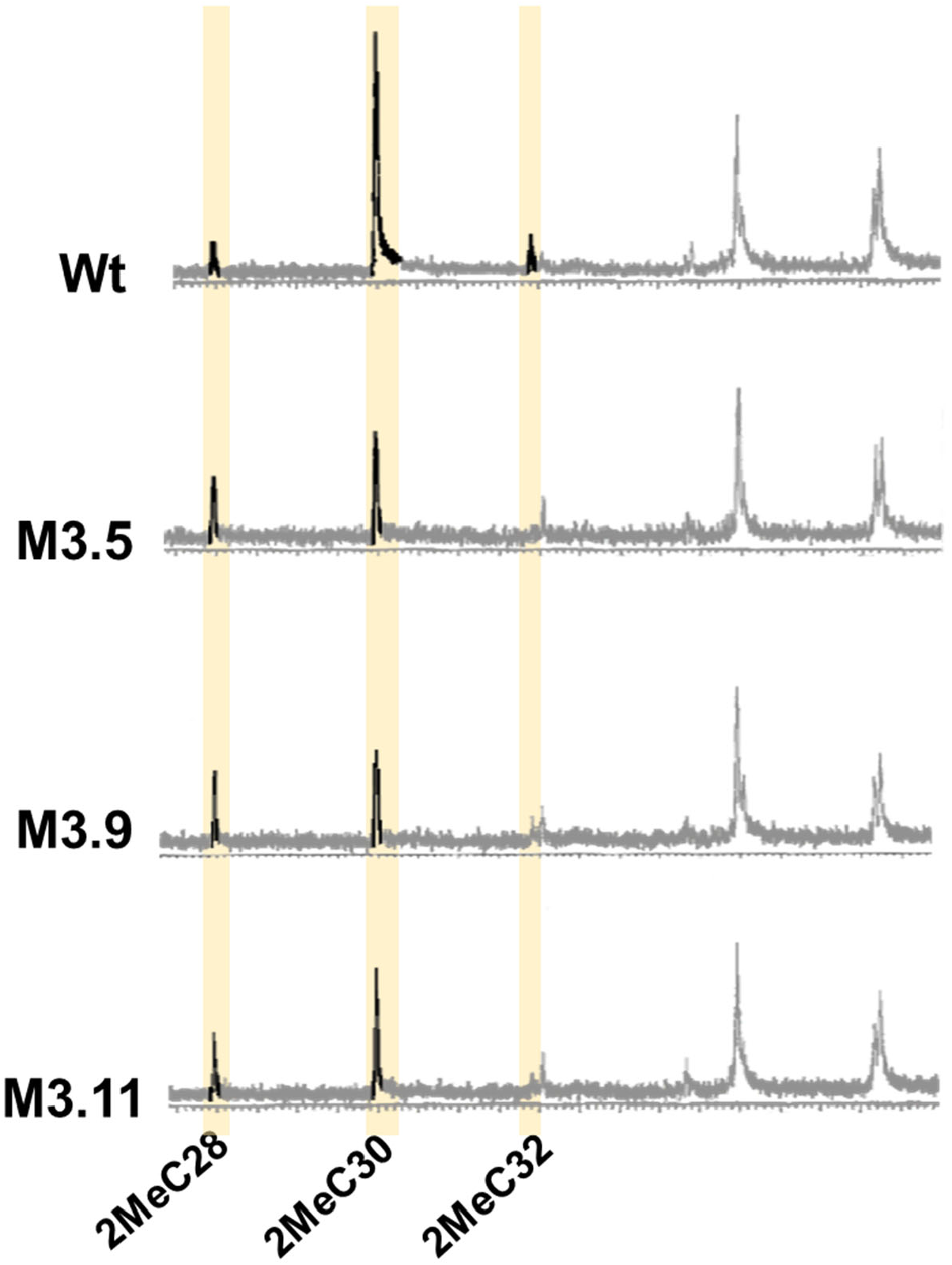
GC-MS chromatograms of mbCHCs in three homozygous *Dmoj/mElo* knockout strains of *D. mojavensis*, M3.5, M3.9 and M3.11. In all three knockout strains, levels of 2MeC30 and 2MeC32 were reduced and levels of 2MeC28 were increased compared to the wild-type control.

**Figure S5.**
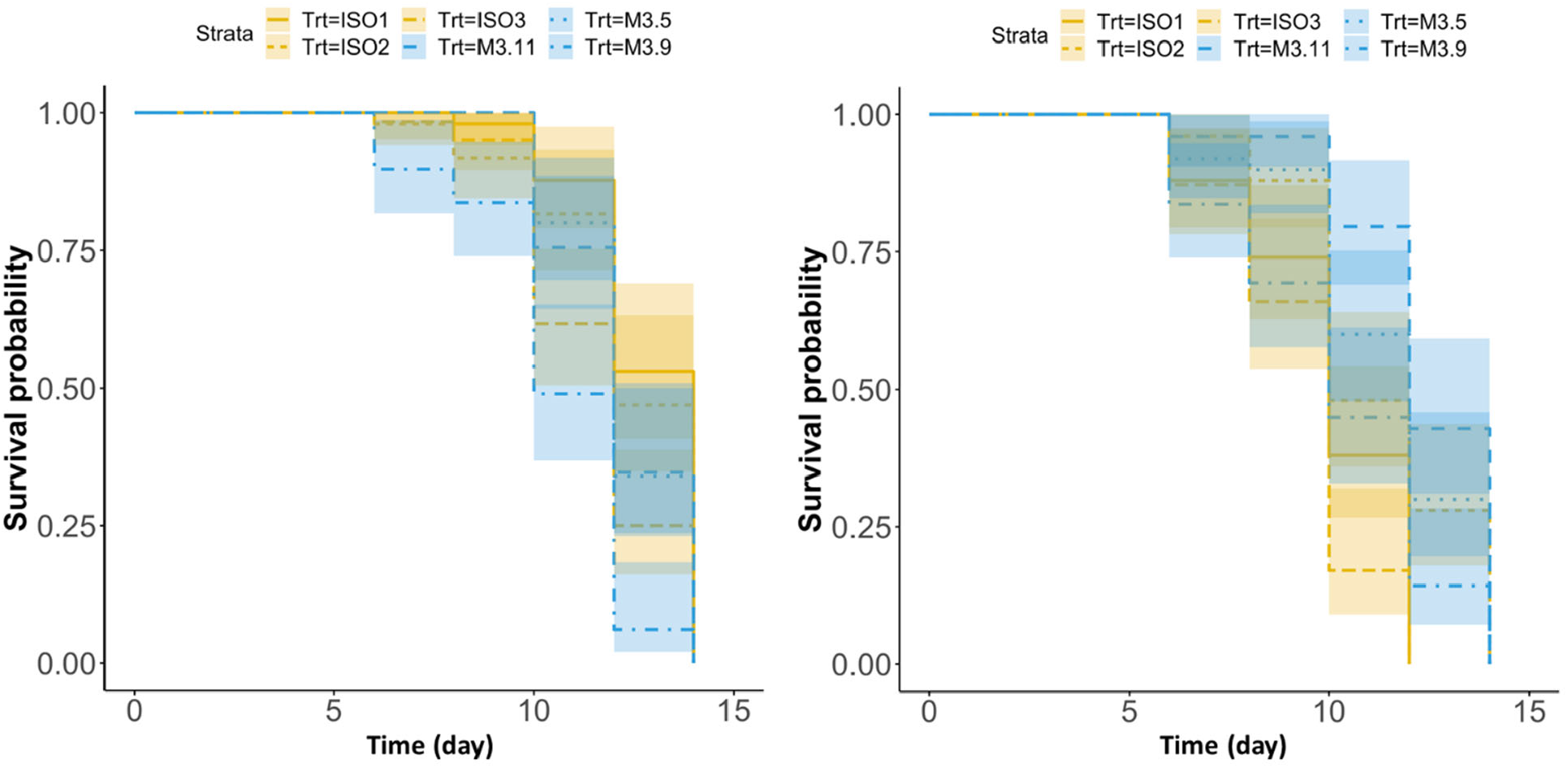
Knockout of the *mElo* orthologue *GI20347* in *D. mojavensis* did not lead to significant differences in survival at 37°C in a non-desiccating environment. Differences in survival between the wild type and *Dmoj/mElo* knockout strains of *D. mojavensis* were determined using linear mixed model with the variation within each group (iso-female or independent knockout strains) being random effects. No significant differences were observed (Female: *P* = 0.4; Male: *P* = 0.2).

**Figure S6.**
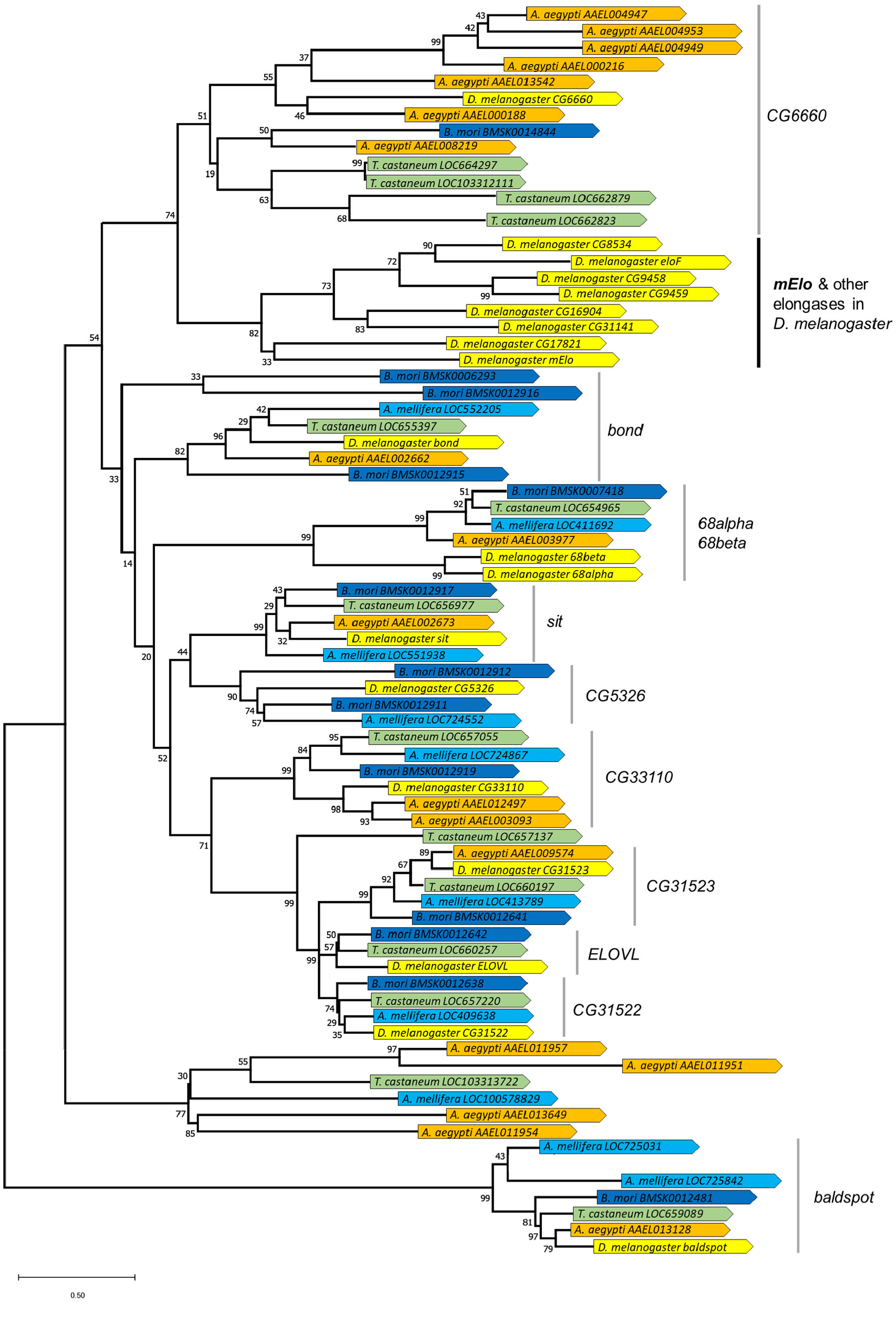
Phylogenetic tree of all elongases genes identified in *Drosophila melanogaster, Aedes aegypti, A*pis *mellifera, Bombyx mori*, and *Tribolium castaneum*. The elongase genes of each species is denoted with a different color. The tree showed that two elongases genes, *CG31523* and *sit*, have one-to-one orthologs across all five species. *mElo* is clustered with a few *Drosophila* genes suggesting that this gene is likely to be *Drosophila* specific.

**Figure S7.**
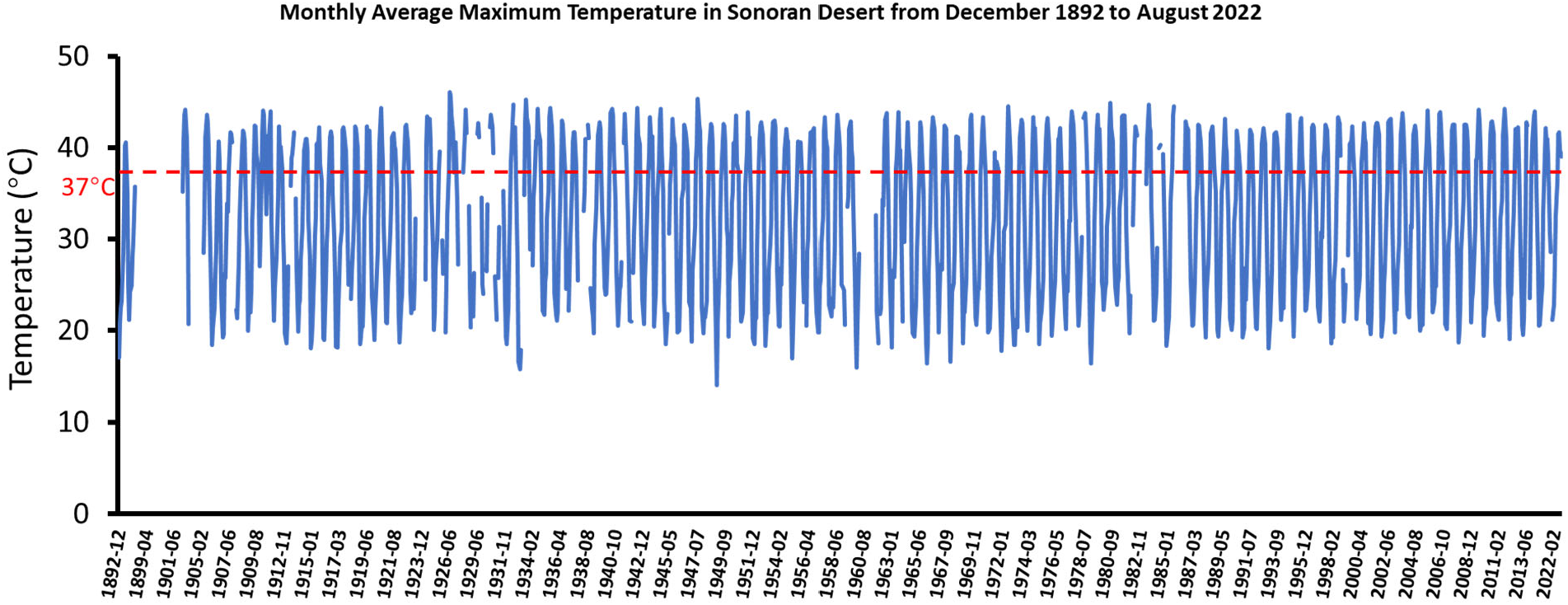
Monthly average maximum temperatures in the Sonoran Desert. Plot of monthly average maximum temperatures in a climatic station (GILA BEND 2 SE, AZ US) in the Sonoran Desert from December 1892 to August 2022. The climatic station is located at the coordinate 32.93803, -112.68109. The red dotted line indicates 37°C. The data were obtained from NCEI-NOAA (https://www.ncei.noaa.gov/).

**Figure S8.**
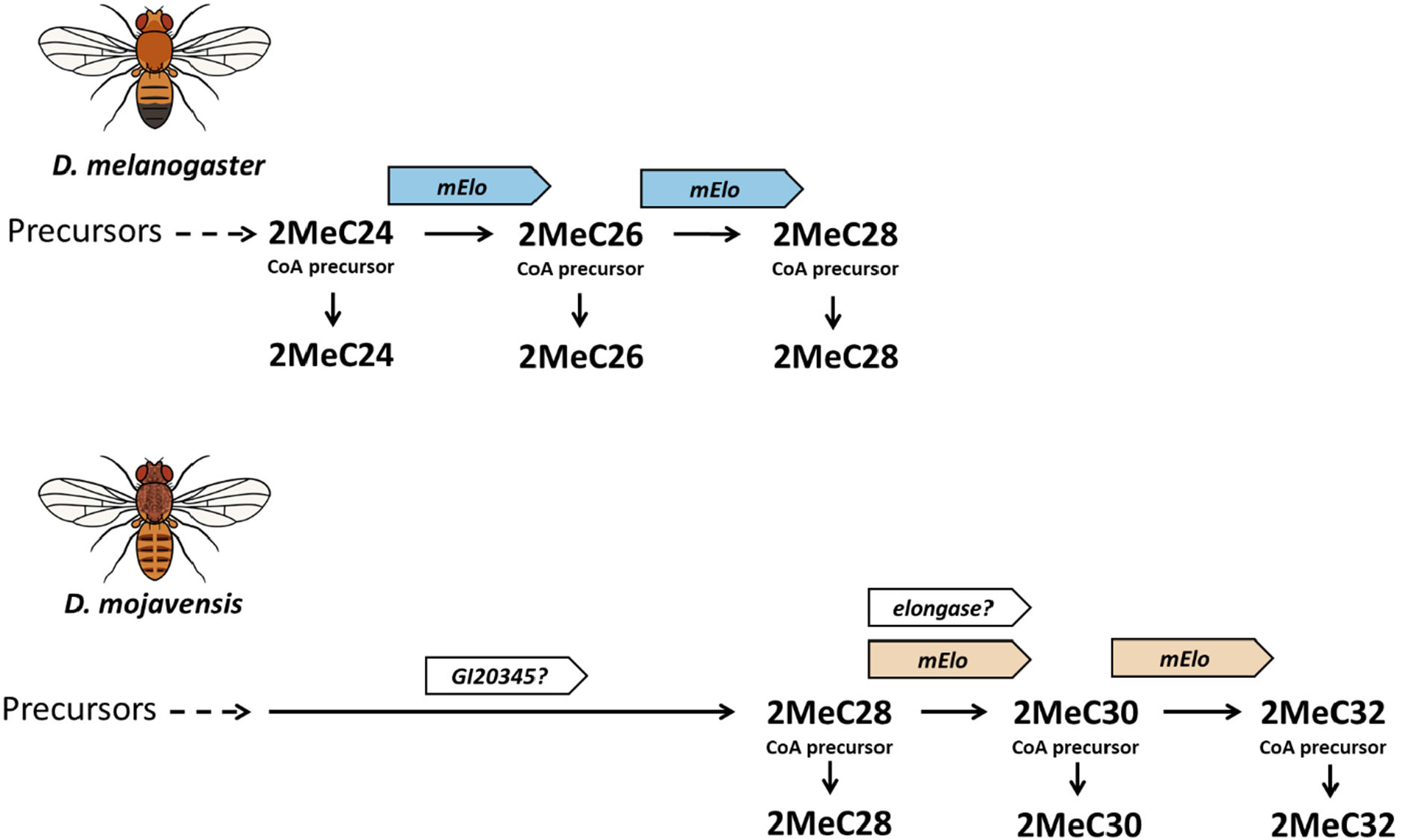
Model showing the production of mbCHCs in *D. melanogaster* and *D. mojavensis*. In *D. melanogaster*, the elongase *mElo* elongates 2MeC24 to 2MeC26 and 2MeC28. In the elongase *mElo* elongates 2MeC28 to 2MeC30 and 2MeC32, while elongation to 2MeC28 is due to another elongase, possibly *GI20345*, which is expressed in *D. mojavensis* oenocytes and can elongate shorter mbCHCs to 2MeC28 when overexpressed in *D. melanogaster* oenocytes. As the knockout of *mElo* did not fully reduce the production of 2MeC30 in *D. mojavensis*, we hypothesize that another elongase may also be involved in the synthesis of mbCHCs up to 2MeC30 in *D. mojavensis*.

**Figure S9.**
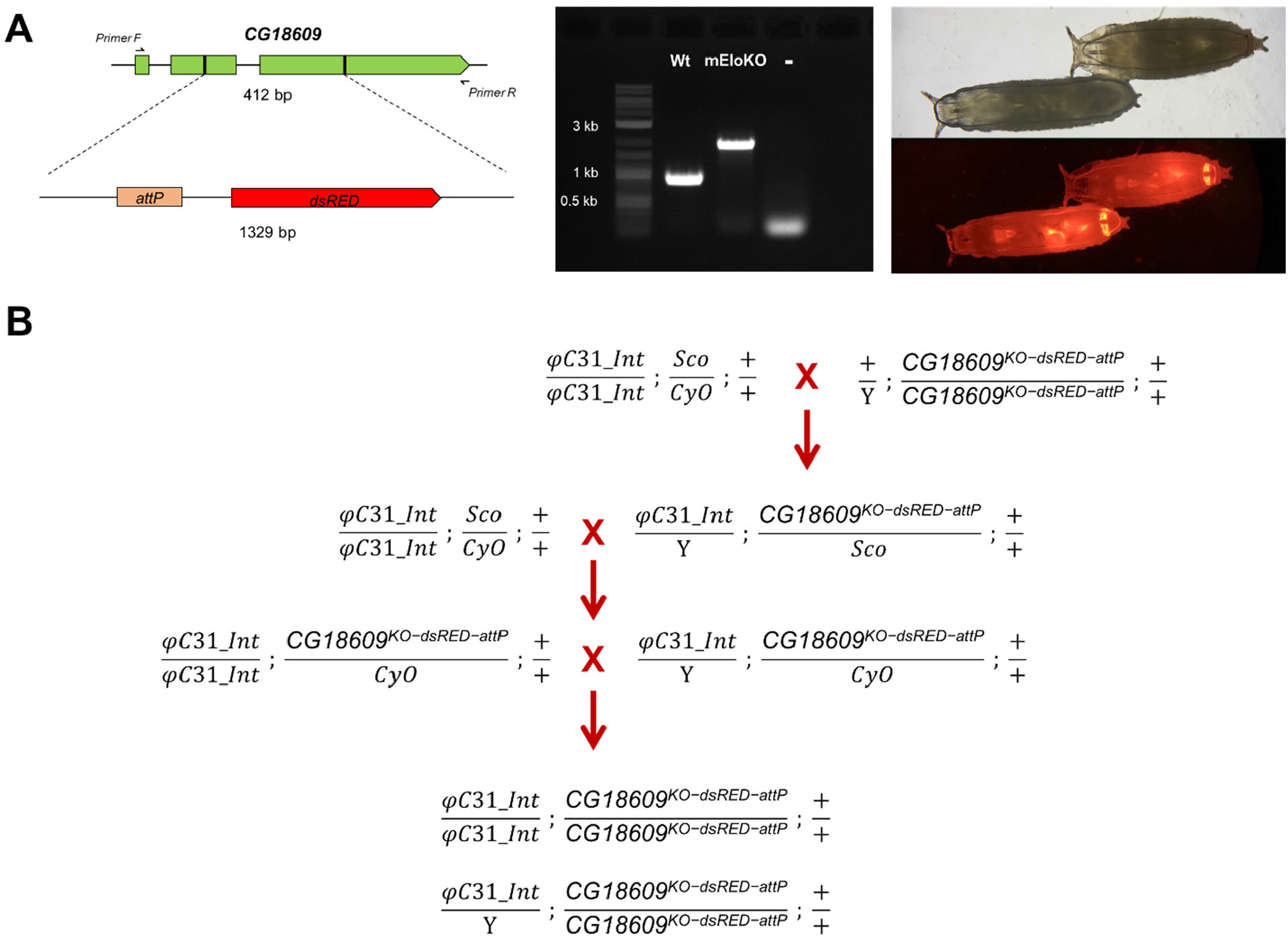
Generation of *mElo* knockout lines in *D. melanogaster*. A. Diagram of knockout using CRISPR/Cas9-mediated homology-directed repair on *Dmel/mElo* (Left panel). Successful knockout was confirmed with the replacement of the *attP*/*dsRED* sequence (Middle panel) as well as the presence of dsRed (Right panel). B. Crossing scheme to generate *w*^*1118*^, *P{y*^*+t7*.*7*^*=nos-phiC31\int*.*NLS}X; CG18609*^*KO-dsRED-attP*^ (*mEloKOint)* strain.

